# Spatiotemporal expression of thyroid hormone transporter MCT8 and *THRA* mRNA in human cerebral organoids recapitulating first trimester cortex development

**DOI:** 10.1101/2023.11.16.567497

**Authors:** Adina Sophie Graffunder, Audrey Amber Julie Bresser, Valeria Fernandez-Vallone, Matthias Megges, Harald Stachelscheid, Peter Kühnen, Robert Opitz

## Abstract

Thyroid hormones (TH) play critical roles during nervous system development and patients carrying coding variants of MCT8 (Monocarboxylate transporter 8) or THRA (Thyroid hormone receptor alpha) present a spectrum of neurological phenotypes resulting from perturbed local TH action during early brain development. Recently, human cerebral organoids (hCOs) emerged as powerful three-dimensional *in vitro* tools for disease modelling recapitulating key aspects of early human cerebral cortex development. To begin exploring prospects of this model for thyroid research, we performed a detailed characterization of the spatiotemporal expression of the TH transporter MCT8 and TH receptor THRA in developing hCOs. Immunostaining of hCOs showed MCT8 membrane expression in neuronal progenitor cell types including early neuroepithelial cells, SOX2+/Nestin+ radial glia cells (RGCs), TBR2+ intermediate progenitors and HOPX+ outer RGCs. In addition, we detected robust MCT8 protein expression in CTIP2+ deep layer neurons and at later developmental stages in SATB2+ upper layer neurons. Spatiotemporal *SLC16A2* mRNA expression, detected by fluorescent *in situ* hybridization (FISH), was highly concordant with MCT8 protein expression across cortical cell layers. FISH detected *THRA* mRNA expression already in neuroepithelium before the onset of neurogenesis and *THRA* expression was maintained in RGCs of the ventricular zone. Increased *THRA* expression was later detected in the subventricular zone whereas highest *THRA* expression was observed in excitatory neurons in peripheral hCO regions. In combination with strong up-regulation of known T3 response genes following short-term T3 treatment of hCOs, these observations show that hCOs provide a promising and experimentally tractable model to probe local TH action during human cortical neurogenesis and eventually to model the consequences of impaired TH function for early cortex development.

## Introduction

Thyroid hormone (TH) signalling is essential for normal brain development and function (1, 2). Studies in different rodent models clearly highlighted a pleiotropic nature of TH action on brain development with regulatory effects of TH on neuronal progenitor dynamics, neuronal migration, layer specification, synaptogenesis, functional maturation, glia cell function and myelination (1, 2). THs exert their function mainly via binding of the biologically active TH 3,3’,5’-triiodothyronine (T3) to nuclear TH receptors (THRA and THRB) that act as ligand-induced transcription factors controlling expression of many genes in neuronal tissues (3–5).

At the target cell level, intracellular amounts of T3 available for TR binding are regulated by complex mechanisms including cellular TH uptake by specialized membrane transport proteins (6). Expression of at least five such transporter proteins (MCT8, MCT10, OATP1C1, LAT1, LAT2), equipped with specific transport capacities for T3 and thyroxine (T4), has been documented in mammalian brain tissues (7, 8). Moreover, cell type-specific transporter expression profiles suggest that different cell types might have distinct capacities for cellular import and export of TH (9–11). Intracellular deiodinases provide another major layer of regulation to control cellular T3 availability (12, 13). Mammalian nervous system tissues show expression of the activating type 2 deiodinase (*DIO2*) and the inactivating type 3 deiodinase (*DIO3*) (14, 15). Reports of *DIO2* expression in astrocytes and *DIO3* expression in neurons again suggest cell type-specific and spatial segregation of TH metabolism in mammalian brain tissue (10).

The essential role of TH signalling for human brain development is reflected by the spectrum of neurological phenotypes of patients experiencing decreased TH availability due to deficient TH synthesis (16, 17) or perturbed local TH signalling caused by inactivating mutations in TH transporters (18, 19) or TH receptors (20, 21). The timing of alterations in TH economy and signalling during development is one critical factor with respect to type and severity of neurological outcomes (22). While rapid initiation of T4 supplementation efficiently rescues neurological development in newborns diagnosed with congenital hypothyroidism (17), therapeutic amelioration of neurological problems can be more limited if perturbed TH function affects brain development throughout gestation (23, 24). One prominent example for the latter situation are patients with Allan-Herndon-Dudley-Syndrome due to mutations in the TH transporter MCT8 (Monocarboxylate transporter 8) (18). These patients present with neurodevelopmental delay including severe psychomotor retardation likely as a consequence of an overall reduced TH availability to developing brain tissue and impaired T3 uptake by neurons and their progenitors (25).

Despite the abundant clinical evidence, there is still a very limited understanding of how tissue-wide TH bioavailability and local TH action are regulated during early human brain development. This knowledge gap is mainly due to inaccessibility of fetal human brain tissue and ethical constraints on experimental work. Accordingly, most of the current knowledge on TH action during mammalian brain development has been derived from rodent models (26–28). However, fundamental inter-species differences in tissue cytoarchitecture, neuronal cell types, developmental features and molecular pathways are complicating human disease modelling in many rodent models (6, 7, 29, 30). The cerebral cortex, for example, has undergone a massive expansion in size and complexity during primate evolution and modelling development of the large, gyrencephalic human cortex based on the small lissencephalic rodent cortex has inherent limitations (29, 31, 32).

The advent of human induced pluripotent stem cell (hiPSC) technologies eventually provided a critically needed resource for *in vitro* modelling of human neurodevelopment and diseases (33). Despite the availability of diverse hiPSC-based protocols to derive cortical neurons and glial cell types, studies on TH action in hiPSC-derived models are still few in number and often limited to two-dimensional cell culture systems (21, 25, 34). In turn, the recent emergence of hiPSC-derived, three-dimensional (3D) human cerebral organoids (hCOs) offers exciting new options to study TH-regulatory action in complex cellular systems closely recapitulating key aspects of human cortex development. These millimetre-sized 3D models provide an unprecedented means to differentiate the full spectrum of neuronal progenitor cell (NPC) populations in spatially segregated domains and to sequentially generate early- and late-born cortical excitatory neurons in an inside-out fashion similar to the human fetal cortex (35–37). The recent past saw the development of different protocols for the generation of cerebral organoids based on self-organizing principles (38–40). It has also been acknowledged that there is variability among different protocols to faithfully generate cortical tissue and that individual hiPSC lines can differ markedly in their organoid-forming capacity (41–44).

With the longer-term goal of establishing cerebral organoids as a robust and sensitive experimental platform for studies on TH function during early human cortex development, a major aim of our study was to benchmark cortical development in organoids generated from in-house hiPSC lines. Based on observations made in culture experiments with different hiPSC lines, we derived a set of critical quality control measures to ensure reproducibility across hiPSC lines and individual organoid batches. We then performed a detailed characterization of the spatiotemporal expression patterns of MCT8 and THRA during key stages of organoid differentiation. Finally, our gene expression analyses demonstrate that hCOs recapitulating first trimester fetal cortex development are responsive to T3 treatment as evident from up-regulation of known TH-responsive genes.

## Material & Methods

### hiPSC maintenance

The healthy donor hiPSC lines BIHi001-B (male), BIHi043-A (female), BIHi005-A (male) and BIHi250-A (female) were cultured on 6 well-plates coated with hESC-qualified Geltrex (Thermo Fisher, #A1413302) in E8 Medium (Thermo Fisher, #A1517001) under hypoxic conditions. Media were changed daily. hiPSC cultures were split upon reaching 80% confluency by incubation in 0.5 mM EDTA diluted in PBS (5 min at 37 °C) and subsequent resuspension of cell clumps in 1.5 ml E8 medium. Cells were seeded with a 1:20 or 1:30 split ratio.

### Cerebral organoids

Cerebral organoids were derived from hiPSC using the STEMdiff™ Cerebral Organoid Kit (StemCell Technologies, #08570, #08571) following the manufactureŕs instructions. Briefly, hiPSC were grown to 60% confluency, washed with PBS and dissociated into a single cell suspension by incubation in TrypLE Select (Thermo Fisher, #12563029) for 5 min at 37°C. Cells were recovered in E8 medium containing 10 µM Rho associated kinase (ROCK) inhibitor Y-27632 (FUJIFILM, #253-00513) and 9,000 live cells were seeded per well of a 96-well Ultra Low Attachment (ULA, Costar, #3473) plate in 100 µl Embryoid Body (EB) Formation Medium. EB cultures were monitored daily under a microscope and 100 µl EB Formation Medium were added per well on days 2 and 4 of culture. Only batches with the large majority of EBs passing general EB quality criteria (round morphology, homogenous cell density) were continued into further culture steps (**quality checkpoint #1**). On day 5, EBs were transferred into Neural Induction Medium and cultured in 24-well ULA plates at a density of four EBs per well. On day 7, high quality EBs were individually embedded into droplets of Geltrex (Thermo Fisher, #A1413202) and further cultured in Expansion Medium in stationary 10 cm cell culture dishes until day 10. At this stage, batches of growing organoids were checked under a microscope for a gross morphological appearance (round morphology, neuroepithelial bud formation) indicative of successful neuroepithelium induction (**quality checkpoint #2**). From day 10 onwards, organoids were cultured in 10 cm cell culture dishes containing 25 to 30 ml Maturation Medium. Organoids were cultured on an orbital shaker at a rotating speed of 65 rpm and media were changed every 2 to 3 days. Between day 24 and day 28 of culture, a subset of organoids from each batch were fixed and analyzed by immunofluorescence (IF) for correct patterning of dorsal pallial tissue (**quality checkpoint #3**). Only those organoids batches that were compliant with all three quality checkpoint criteria were continued for culture up to 10 weeks.

### Cryosectioning

At indicated time points, organoids were randomly sampled into ice-cold 1X PBS solution and subsequently fixed in 4% paraformaldehyde (PFA) solution in 1X PBS overnight at 4°C with gentle agitation. After several 10 min washes in 1X PBS, organoids were immersed overnight in 30% sucrose in 1X PBS and embedded in Tissue-Tek 4566 cryomolds (Thermo Fisher, # 10844231) using optimal cutting temperature (OCT) embedding matrix (Carl Roth, #6478). Cryomolds were placed on dry ice and fixed frozen tissue blocks were stored at −20°C before sectioning. Tissue blocks were sectioned at 20 - 30 µm thickness onto SuperFrost Plus microscope slides (Fisher Scientific, #J1810AMNZ) using a CryoStar NX70 (Thermo Fisher) cryostat.

### Immunofluorescence staining

For IF staining, tissue sections were air-dried for at least 30 min and if indicated (see **Supplementary Table S1**), citrate antigen retrieval (10 mM sodium citrate, pH 5.8) was performed by boiling tissue sections in a microwave for 7 min. Sections were washed several times in 1X PBS to remove residual OCT and were then immersed in 1X blocking solution, consisting of 5% normal donkey serum (Jackson ImmunoResearch, #017-000-121) and 0.3% Triton-X (Sigma, #T8787) in 1X PBS, for 60 min at room temperature. Primary antibody incubation in 0.1X blocking solution was performed overnight at 4°C. Sections were washed three times for 10 min with 1X PBS and incubated with secondary antibodies in 0.1X blocking solution for 2 h at room temperature. If indicated, staining of F-actin using Phalloidin (abcam, #ab176759) was combined with the secondary antibody incubation step. A list of all antibodies used in this study is provided in **Supplementary Table S1**. Sections were washed three times for 10 min with 1X PBS and incubated in 4,6-diamidino-2-phenylindole (DAPI, 1 µg/mL) solution to counterstain cell nuclei. Sections were coverslipped using Roti®-Mount FluorCare mounting medium (Carl Roth, #HP19.1).

### RNAscope fluorescence *in situ* hybridization

We used the RNAscope Multiplex Fluorescence Assay v1 (Advanced Cell Diagnostics, #320850) to examine spatial expression profiles of *SLC16A2* mRNA (RNAscope Probe - Hs-SLC16A2, #562191) and *THRA* mRNA (RNAscope Probe - Hs-THRA-C2, #562201-C2). All RNAscope experiments were combined with IF detection of at least one informative cell type marker (i.e. SOX2, TBR1) using the Co-Detection workflow according to the manufacturer recommendations. Briefly, sections were initially processed as described in the IF staining protocol and incubated in primary antibodies overnight at 4°C. The main modification here was the use of Co-Detection Antibody Diluent (Advanced Cell Diagnostics, #323160) for blocking and antibody incubation steps. After removal of primary antibodies, sections were washed several times in 1X PBS and post-fixed in 4% PFA for 10 min at room temperature. After further washes in 1X PBS, sections were gradually dehydrated in 50%, 70%, 100% and 100% ethanol and incubated in protease IV solution (Advanced Cell Diagnostics, #322336) for 30 min at room temperature. The following *in situ* hybridization steps (pooled probe hybridization and signal amplification) were performed at 40°C according to the manufacturer’s protocols. After completion of the RNAscope hybridization protocol, sections were washes two times 5 min in 1X PBS and incubated with secondary antibodies in Co-Detection Antibody Diluent overnight at 4°C. Counterstaining of cell nuclei and coverslipping was done as described above. RNAscope experiments also included hybridization of tissue sections with RNAscope 3-plex negative control probes and RNAscope 3-plex positive control probes (Advanced Cell Diagnostics, #320871 and #320861). Positive control probes were targeting *POLR2A* in channel C1, *PPIB* in channel C2, and *UBC* in channel C3. Negative control probes were targeting bacterial transcripts.

### Confocal imaging

Fluorescent images of IF- and RNAscope-stained sections were acquired on a Leica TCS SP8 confocal microscope with x25 and x63 water-immersion objectives using LAS-X software (Leica). Tile scan images of whole organoid sections were captured using a 25× water-immersion objective and stitched using LAS-X Software. RNAscope-stained sections were imaged as a *z*-stack of five images (1 µm distance) and maximum intensity projection views were generated using LAS-X software.

### T3 treatment of organoids

Organoids derived from three hiPSC lines (BIHi001-B, BIHi043-A, BIHi250-A) were cultured in parallel under standard conditions described above until day 44 of culture. At this time point, 10 organoids per hiPSC line were allocated to 6 cm cell culture dishes containing either standard medium (control group) or standard medium supplemented with 50 nM T3 (T3 treatment group). Organoids were then cultured for a total of 48 h on an orbital shaker at a rotating speed of 65 rpm without any additional media exchanges. At the end of the 48 h treatment period, pools of 2-3 organoids were flash frozen in liquid nitrogen and stored at −80°C until RNA extraction.

### RNA isolation and real-time quantitative PCR

Total RNA was isolated from pools of 2-3 organoids using RNeasy Plus Mini Kit (Qiagen, #74134) according to the manufacturer’s protocols. RNA concentrations were measured on a NanoDrop One Spectrophotometer (Thermo Fisher). Per sample, 200 ng of total RNA was reverse transcribed using Oligo(dT)_20_ Primer (Thermo Fisher, #18418020) and SuperScript III Reverse Transcriptase (Thermo Fisher, #18080044). Real time-qPCR was carried out in 10 µl reactions on MicroAmp EnduraPlate 384-well plates (Thermo Fisher, #4483321) using SYBR® Green PCR Master Mix (Thermo Fischer, #4309155). Real-time PCR was performed on a QuantStudio 6 Real-Time PCR System (Thermo Fisher). Primers used in Real time-qPCR experiments are listed in **Supplementary Table S2**. All reactions were performed in duplicates. Relative quantifications of target transcripts were done based on the 2^-ddCT^ method using *UBE2D2* and *TBP* as control genes.

## Results

Approaches for forebrain organoid generation differ in complexity of protocols, technical equipment requirements, use of guidance factors and extracellular matrix components, organoid size, representation of forebrain regions as well as temporal dynamics of cell differentiation (45–48). In this study, we generated cerebral organoids using a commercially available kit of reagents that supports an organoid culture strategy initially proposed by Lancaster and Knoblich (46). Key steps of our organoid generation protocol are highlighted in **Fig. 1A** and changes in gross morphological characteristics of developing organoids are shown in **Fig. 1B**. Cerebral organoids generated in this study contain a variable number of small cortical units. These units are composed of a neural rosette-like inner part surrounded by specific neuronal cell populations that are arranged in a laminar fashion (see **Fig. 1C**). Over the course of 10 weeks, these organoids grow to a size of approximately 3 mm in diameter. A representative growth trajectory of organoids generated with the BIHi001-B line is shown in **Fig. 1D**.

**Figure 1.**
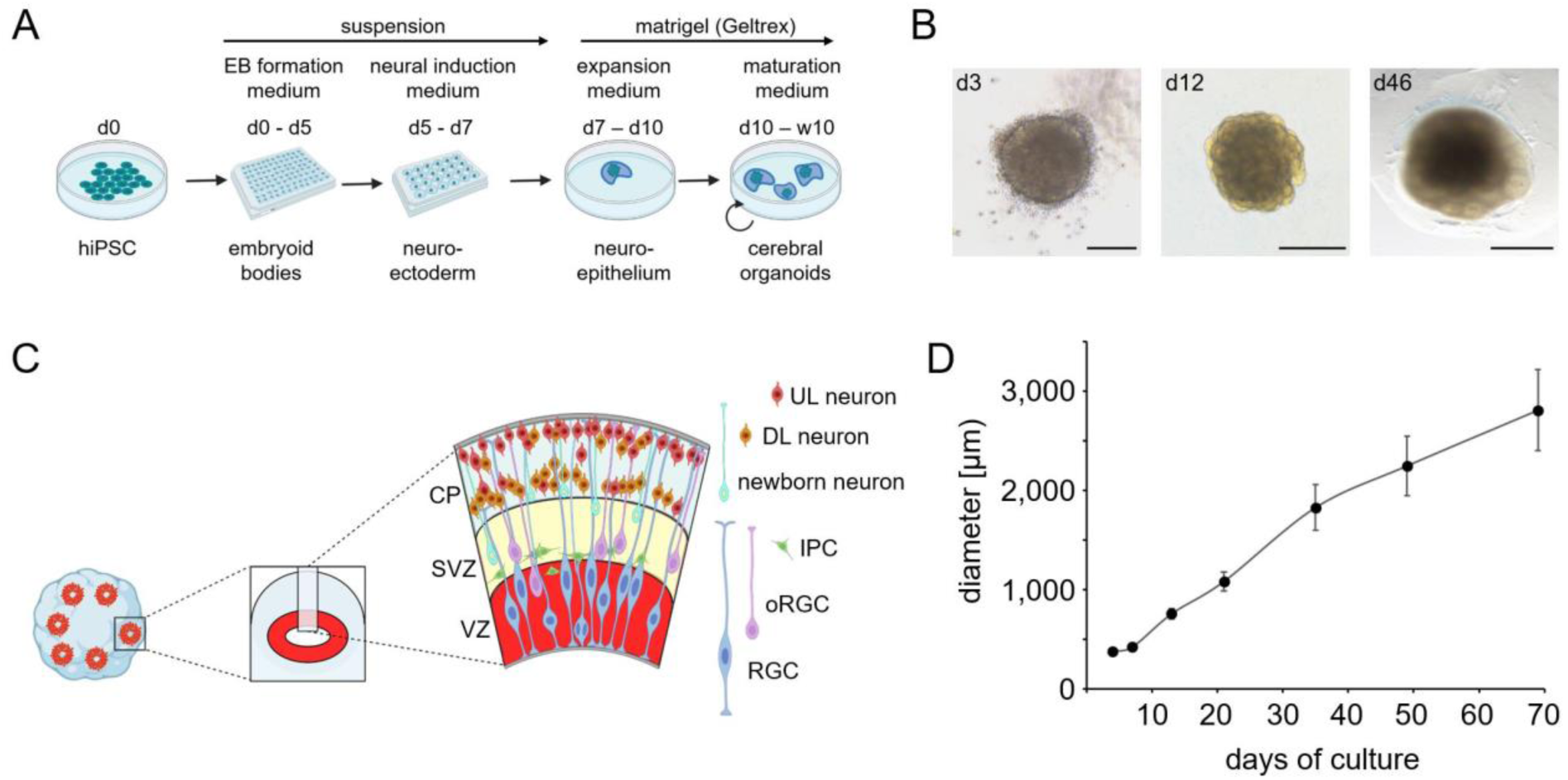
(**A**) Schematic overview of key steps of the protocol used to derive cerebral organoids from hiPSC. The culture protocol promotes generation of organoids recapitulating dorsal pallial tissue differentiation. (**B**) Brightfield microscopic images of organoids at three stages of differentiation including EB stage on day 3 (scale bar: 200 µm), neuroepithelial stage on day 12 (scale bar: 500 µm) and neurogenesis stage on day 46 of culture (scale bar: 1 mm). (**C**) Scheme summarizing the laminar organization of cell types in 9-week old organoids. The ventricular zone (VZ) hosts radial glia cells (RGC), the main neural stem cell pool in cerebral organoids. Asymmetric divisions of RGC at the apical pole of the VZ yield intermediate progenitor cells (IPC) and outer RGC (oRGC) which migrate out of the VZ and populate apical (IPC) or more basal regions (oRGC) of the subventricular zone (SVZ). Newborn neurons generated by IPC and oRGC in the SVZ migrate to the cortical plate (CP) where early-born, deep layer (DL) neurons populate inner CP regions and later-born, upper layer (UL) neurons are enriched in the most peripheral CP regions. (**D**) Growth trajectory of organoids derived from BIHi001-B line during a 10-week culture experiment. Measurements of the diameter (in µm) of the cross-sectional area of phase contrast images are shown. Data presented are means ± standard deviations (N=10-15 organoids per time point). Panels **A** and **C** were created with BioRender.com.

### Cytoarchitecture of developing cerebral organoids

Because a review of the available literature on growth and differentiation dynamics in different organoid-based studies showed a remarkable variability, we first performed a series of experiments with the aim to benchmark the temporal sequence of neuronal development in organoids cultured under our in-house conditions. Immunostainings shown in **Fig. 2** illustrate developmental changes in gross cytoarchitectural organization of cerebral organoids at key stages of organoid differentiation. Early-stage organoids collected at 2 weeks of culture were almost entirely composed of neuronal progenitor cells (NPC) co-expressing the neuroepithelial markers SOX2 and Nestin (**Fig. 2A-F**). Uniform expression of the telencephalic marker FOXG1 (**Fig. 2B** and **Supplementary Fig. S1**) and the dorsal pallial marker PAX6 (**Supplementary Fig. S1**) in SOX2+ NPC confirmed robust specification of a cortical NPC population that will serve as neural stem cells of the cortical excitatory neuron lineage. Markers of newborn neurons (DCX, TUJ1) were not yet detectable at this early stage of our protocol (data not shown). Tissue sections of 2-week-old organoids showed that SOX2+ NPC undergo progressive organization into rosette-like structures (see **Fig. 2E**). Size of rosettes and luminal cavities were still variable for any given tissue section at this early developmental stage. We also noted that 2-week-old organoids contained areas where NPC are not yet organized in rosette-like structures (see **Fig. 3A-F**).

**Figure 2.**
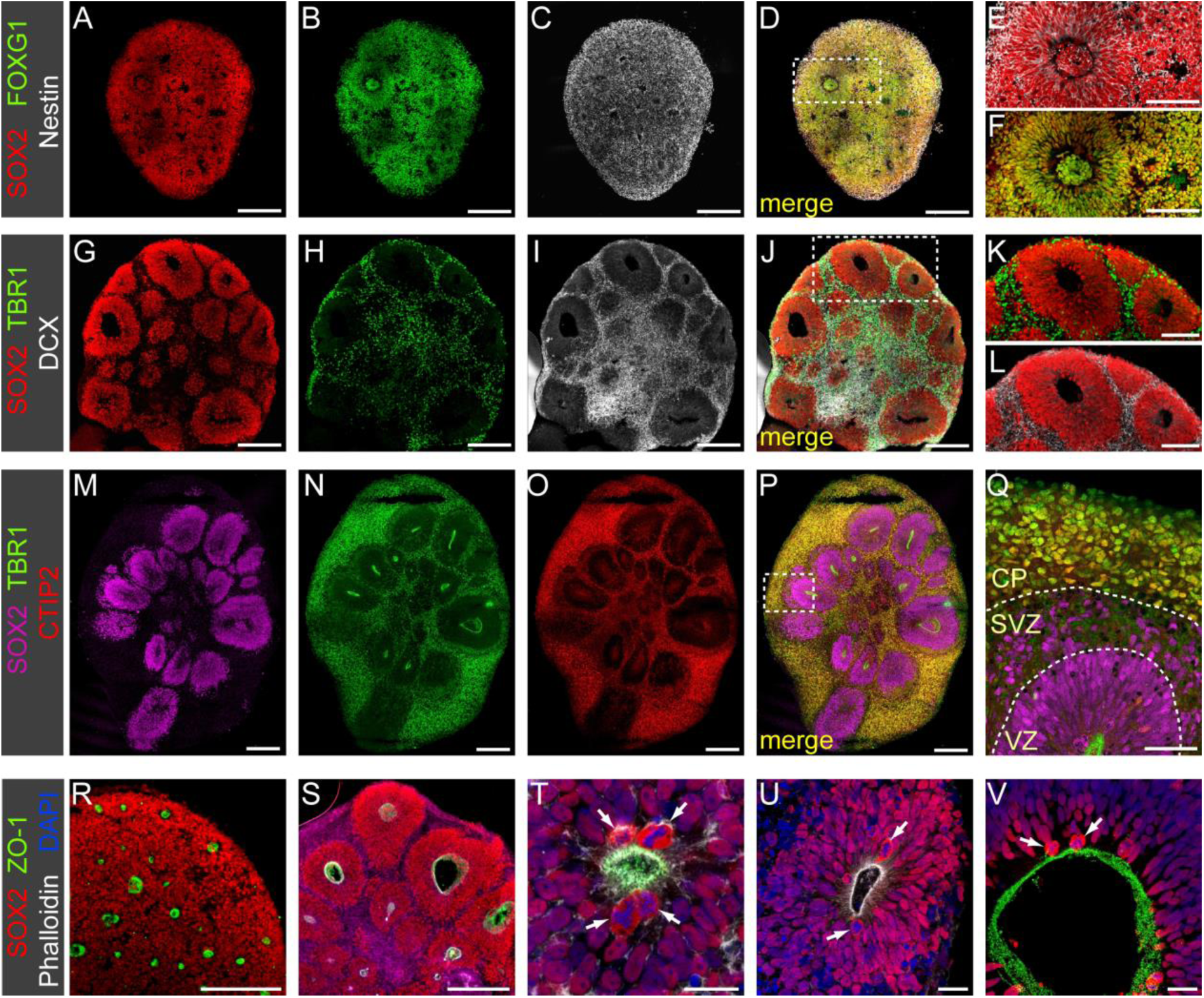
Confocal images of immunostained sections illustrating developmental changes in cytoarchitecture of organoids. (**A-F**) 2-week-old organoids were composed of SOX2+/Nestin+ neuronal progenitor cells (NPC) expressing the forebrain marker FOXG1. Formation of first few neuronal rosettes was evident at this stage (panels **E** and **F** show magnified views of the boxed region in **D**). Markers of neurons were absent at this stage. (**G-L**) 4-week-old organoids were in early stages of neurogenesis. The now larger rosettes are composed of a pseudostratified neuroepithelium of SOX2+ NPC surrounding a central lumen. Newborn neurons expressing doublecortin (DCX) and the cortical neuron marker TBR1 surround individual rosettes, often as a thin layer separating individual rosettes (panels **K** and **L** show magnified views of the boxed region in **J**). (**M-Q**) In 6.5-week-old organoids, the number of TBR1+ neurons is greatly increased and a cortical plate-like zone (CP) has formed in peripheral regions of organoids. At this stage, the majority of TBR1+ neurons co-express the deep layer neuron marker CTIP2. Formation of a distinct subventricular zone (SVZ) separating the ventricular zone (VZ) and the CP is evident. Note the abundant TBR1^low^ newborn neurons located next to SOX2+ progenitors within the SVZ (panel **Q** shows magnified view of the boxed region in **P**). Dashed lines in **Q** mark the border between VZ, SVZ and CP. (**R-V**) Rosettes show hallmarks of a polarized neuroepithelium. The luminal border of the VZ is labelled with apical domain markers (ZO-1, Phalloidin-stained F-actin) throughout organoid differentiation. Images show sections of 2-week-(**R, T**), 4-week-(**S, U**) and 7-week-old organoids (**V**). Apico-basal polarity of the neuroepithelium is also reflected by restriction of cell divisions (arrows in **T**, **U**, **V**) to the apical border. Scale bars: 200 µm (**A-D, G-J, M-P, S, R**), 100 µm (**E, F, K, L**), 50 µm (**Q**), 20 µm (**U-V**).

**Figure 3.**
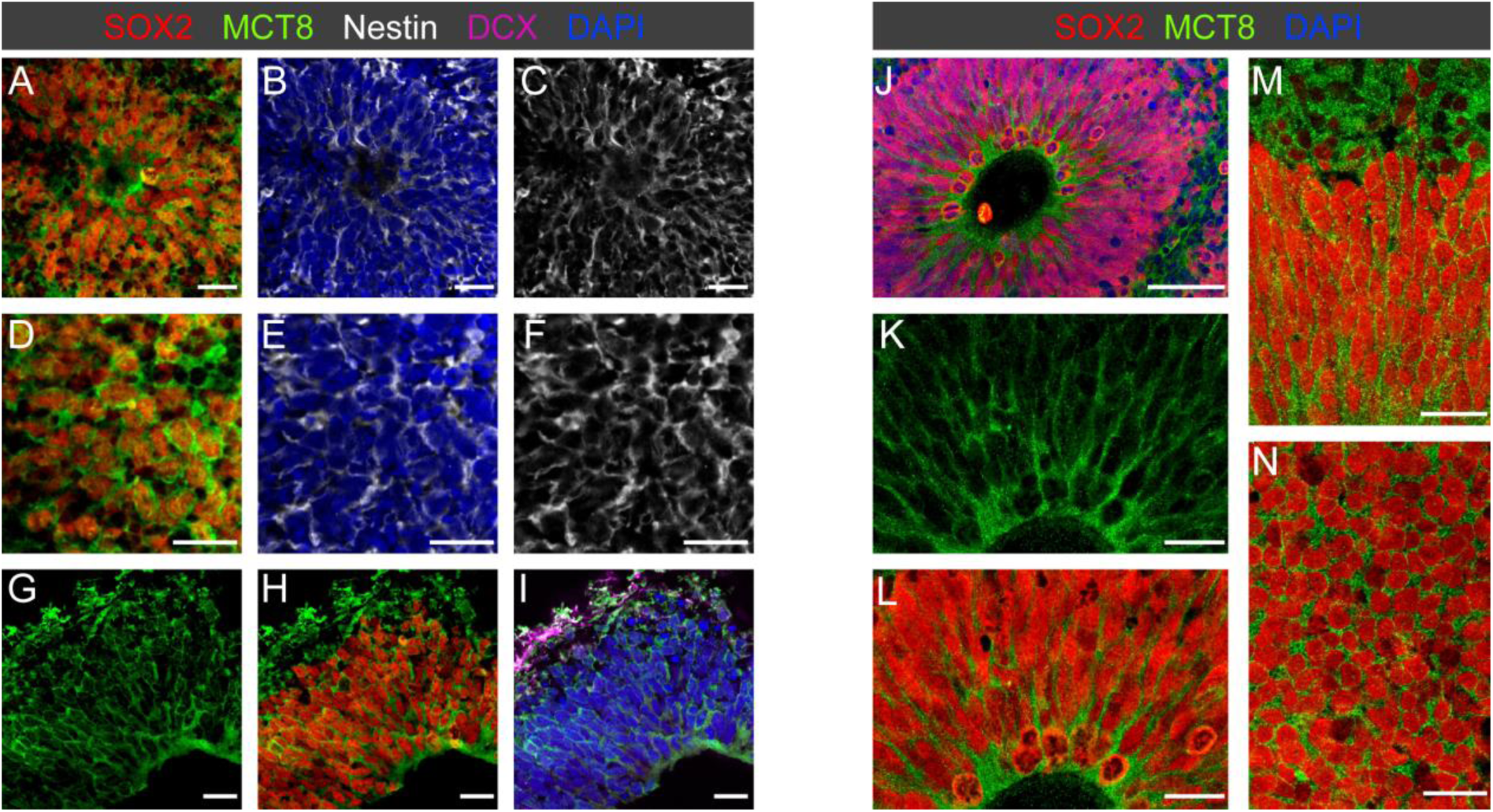
MCT8 is expressed in early neuroepithelium. (**A-F**) In 2-week-old organoids, membrane MCT8 staining is prominent in SOX2+/Nestin+ neuronal progenitor cells (NPC). Note that MCT8 expression was present in both NPCs organized in rosettes (**A-C**) and NPCs not yet arranged in a structured fashion (**D-F**). (**G-H**) At the onset of neurogenesis in 3-week-old organoids, membrane MCT8 staining was observed in SOX2+ NPCs in the neuroepithelium and the first few DCX+ newborn neurons. Note the enhanced MCT8 staining signal at the apical border of the neuroepithelium facing the ventricular lumen. (**J-N**) Mitotic RGC at the apical pole of the VZ of 4.5-week-old organoids maintain MCT8 expression. Analysis of different sectioning planes throughout the VZ (see **M, N**) shows a homogenous MCT8 immunostaining intensity of RGC. Scale bars: 50 µm (**J**), 20 µm (**A-I, K-N**).

Rosette and apical lumen size increased from week 2 to week 4 of organoid culture (**Fig. 2G**). The concomitant increase in organoid size during this period can be attributed in a large part to an expansion of the NPC population. By week 3, the neuroepithelium of all rosettes had assumed a pseudostratified organization to accommodate the increasing number of radial glia cell (RGC) bodies along the apico-basal axis (**Fig. 3H**). Based on cytoarchitectural hallmarks, the neuroepithelium of rosettes in 3-week-old organoids resembles the ventricular zone (VZ) of the *in vivo* cortex at about gestational week 6.5 (49) and we refer to this germinal zone of organoids as VZ from this stage onwards. First few post-mitotic neurons were detectable in 3-week-old organoids and such newborn DCX+ neurons were located adjacent to the basal VZ border (see **Fig. 3I**). These earliest neurons were likely generated by direct neurogenesis from RGC (**Supplementary Fig. S3**) as previously reported in fetal cortex and *in vitro* grown hCOs (49, 50).

By four weeks of culture, organoids presented a distinct morphological organization characterized by enlarged rosettes surrounded by a thin layer of TBR1+/DCX+ neurons (**Fig. 2G-L**). This stage of organoid differentiation is also characterized by the appearance of first TBR2+ intermediate progenitor cells (IPCs) at the basal side of the VZ marking the switch from direct to indirect neurogenesis during the fourth week of organoid culture. A corresponding cytoarchitectural organization has been reported in human embryos at about gestational weeks 7.5 (49).

Between week 4 and 7 of culture, organoids developed multi-layer stratified structures resembling in many molecular, neuroanatomical aspects the *in vivo* morphogenesis of the cortical wall between gestational weeks 8 and 12 (49, 51). Most prominently, there is a massive increase in the total number of TBR1+ cortical neurons particularly in peripheral regions of organoids (compare **Fig. 2H** and **Fig. 2N**). At 5 weeks of culture, the majority of these TBR1+ neurons co-express the deep layer neuron marker CTIP2 (**Fig. 2N-Q**). Expression of the upper layer neuron marker SATB2 was detectable from 8 weeks onwards (see **Fig. 5**). Thus, cerebral organoids recapitulated the temporal order of *in vivo* cortical neuron generation by first producing neurons with deep layer identity (CTIP2+) before switching later to the production of upper layer neurons (52). Higher magnification views of individual rosettes in 6.5-week-old organoids (see **Fig. 2Q**) show the formation of three major laminar zones including a VZ containing densely packed SOX2+ RGC, a subventricular zone (SVZ) containing different populations of NPC intermixed with newborn TBR1^low^ neurons and a peripheral cortical plate (CP) zone populated exclusively by neurons expressing high levels of TBR1 and CTIP2. This laminar neuroanatomical organisation of organoid tissue including VZ, SVZ and CP was maintained until 10 to 12 weeks of organoid culture.

Apico-basal polarity is a key feature of the developing neuroepithelium lining the ventricles *in vivo* (53). Staining of organoid tissue for markers of the apical surface of polarized epithelia including ZO-1 (tight junctions) and phalloidin (F-actin assembly) (**Fig. 2R-V**) demonstrated apico-basal polarity not only in larger rosettes (**Fig. 2U, V**) but also revealed many small cavity spots indicating the formation of numerous nascent neuronal rosettes in early-stage organoids that are otherwise difficult to discern (**Fig. 2R-T**). Another prominent feature of neuronal rosettes was that cell divisions were almost exclusively present at their apical domains (see arrows in **Fig. 2T-V**). This observation is consistent with descriptions of the neuroepithelium in other cerebral organoid studies and in the fetal human cortex (54, 55). The translocation of nuclei for mitosis at the apical surface (known as interkinetic nuclear migration) reportedly occurs similarly in *in vitro* grown cerebral organoids and *in vivo* embryonic forebrain (54).

### Quality control measures

When comparing organoids derived from different healthy donor hiPSC lines, we noticed very similar time courses of neuronal development for lines BIHi001-B (male), BIHi250-A (female) and BIHi043-A (female) confirming the reproducibility of our culture approach across different hiPSC lines. However, in experiments with another hiPSC line, BIHi005-A (male), we observed a much more variable outcome of organoid differentiation. Organoids derived from the latter line often lacked dorsal pallial patterning at early stages of differentiation (see **Supplementary Fig. S2**). Based on these observations and in order to ensure reproducibility and robustness across individual batches of organoids, we devised an organoid batch quality assessment strategy. For this purpose, we defined three quality control checkpoints during early stages of organoid differentiation based on gross morphological characteristics of developing organoids in combination with immunostaining of markers (FOXG1, PAX6, TBR1) indicative of successful dorsal pallial patterning in early-stage organoids (for details see Material and Methods section). Only those organoid batches that were compliant with quality control criteria were included in the following analysis of MCT8 and THRA expression profiles during organoid differentiation.

### MCT8 expression in the developing VZ of cerebral organoids

The severe neurological phenotypes presented by patients with MCT8 mutations have generated great interests in understanding the neurodevelopmental mechanisms underlying this disorder. Given that knowledge on the spatiotemporal expression of this critical TH transporter during early human cortex development is still fragmentary (10, 56, 57), we performed immunostainings of MCT8 in a large developmental series of organoids collected between week 2 and 10 of organoid culture. The MCT8 antibody (RRID:AB_1079343) used in our experiments has recently been used to characterize MCT8 expression in human fetal cortex tissue (10).

The earliest organoid tissue samples analyzed were from 2-week-old organoids. Immunostainings showed a strong MCT8 membrane expression of SOX2+/Nestin+ cells, irrespective of whether NPCs were organized in rosette-like structures (**Fig. 3A-C**) or were still arranged in a less structured fashion (**Fig. 3D-F**). By 3 weeks of culture, MCT8 expression was detected throughout the VZ-like thickened, pseudostratified neuroepithelium as well as in the sparse DCX+ newborn neurons located adjacent to the VZ (**Fig. 3G-I**). While the intensity of MCT8 immunolabelling of cell bodies of SOX2+ RGC was homogenous throughout the thickness of the VZ layer, we observed an enhanced staining intensity at the apical neuroepithelial surface. Such enhanced MCT8 immunolabelling of RGC apical end-feet close to the luminal cavity was even more prominent in the expanded VZ of 4-week-old organoids (**Fig. 3J-L**). The laminar cytoarchitecture of 4-week-old organoids resembles very early stages of *in vivo* cortical development (GW 7.5 to 8) and it is therefore notable that examination of human fetal cortex at later mid-gestation stages also reported that MCT8 staining in the VZ was stronger at the apical surface of the VZ (10).

The apical surface of the VZ was lined by numerous dividing cells (see **Fig. 3L**) and high magnification microscopy showed that MCT8 membrane expression was also detectable in dividing cells see (**Fig. 3K, L**). MCT8 immunostaining of the VZ in different sectioning planes showed a homogenous MCT8 immunostaining intensity across the population of SOX2+ RGC (**Fig. 3M, N**). This pattern of MCT8 expression in the VZ was very much unchanged throughout all later stages of organoid development (up to 10 weeks of culture) and corresponds to the VZ expression pattern previously reported for second trimester fetal cortex tissue (10).

### MCT8 expression in the SVZ of cerebral organoids

Formation of a morphologically distinct SVZ-like germinal layer started at about 4.5 weeks of organoid culture. At this stage of development, asymmetric divisions of RGC in the VZ gave rise to TBR2+ IPCs that delaminated from the VZ and populated the region just basal to the VZ (**Fig. 4A-E**). In contrast to RGC, IPC lack apical or basal processes and a defined cell polarity (49, 58). Basally positioned TBR2+ IPC showed reduced or no SOX2 expression at all in our IF stainings of organoid section (**Fig. 4D, E**). In the fetal cortex, basally located IPCs usually undergo a limited number of amplifying symmetric divisions before undergoing a final self-consuming symmetric neurogenic division (59). Because of the primary role of IPCs for early neurogenesis during these early stages, we analyzed MCT8 expression in IPCs of 4- to 6-week-old organoids. Irrespective of their specific position within the VZ or the SVZ, TBR2+ IPCs always showed a strong membrane MCT8 staining that was similar or slightly stronger compared to neighbouring SOX2+ RGC (**Fig. 4F-J**).

**Figure 4.**
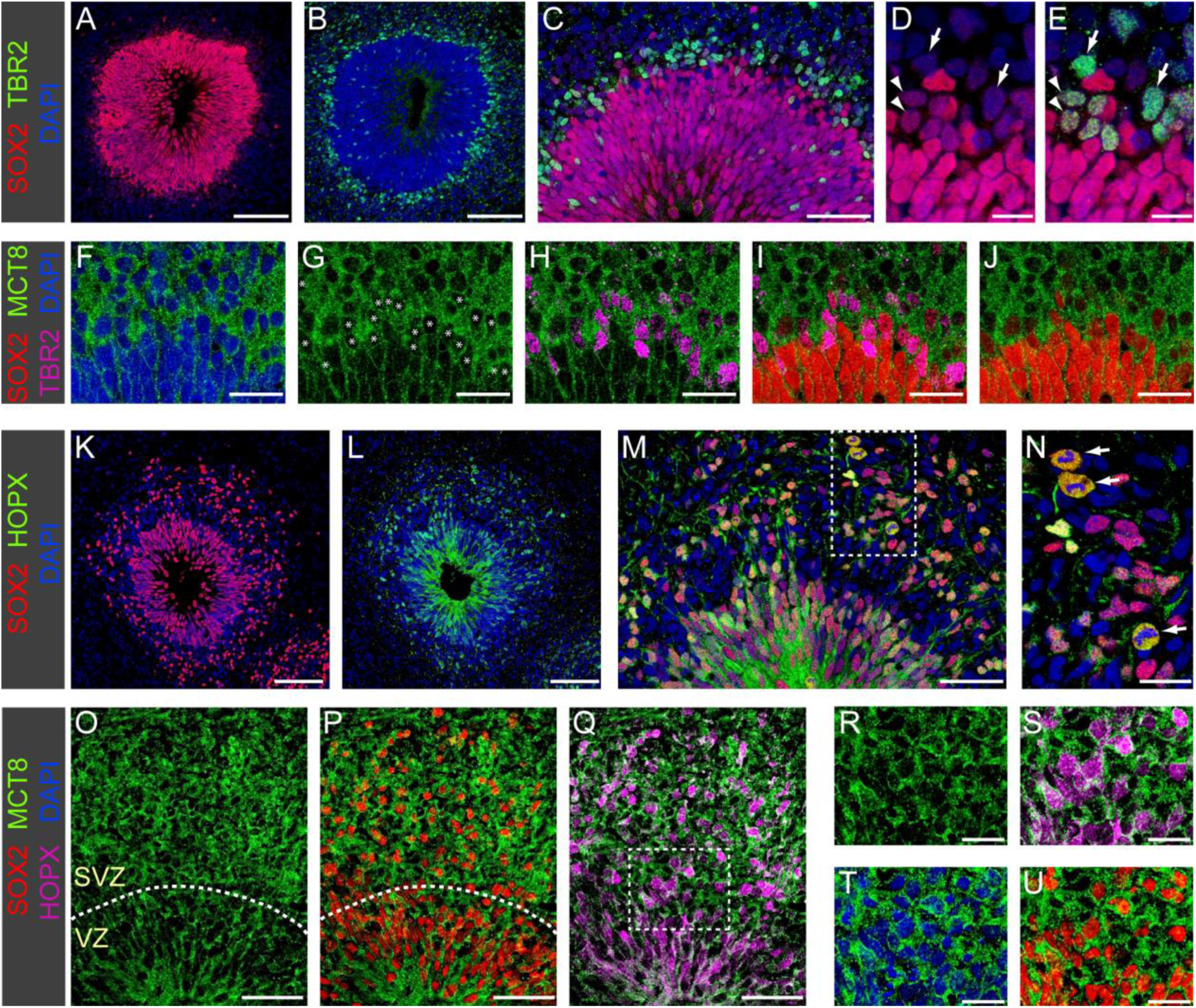
MCT8 expression in neurogenic progenitors. (**A-E**) Formation of a distinct cell layer containing TBR2+ IPC basal to the VZ in 4.5-week-old organoids. IPC generated by asymmetric RGC divisions in the apical VZ migrate towards more basal positions (see **C** for migrating TBR2+ IPC within VZ) and IPC located at the basal VZ border show either low SOX2 (arrowheads in **D, E**) or no SOX2 expression (arrows in **D, E**). (**F-J**) Immunostaining of 5-week-old organoid shows strong MCT8 expression in IPCs. Asterisks in **G** mark the cell bodies of TBR2+ IPCs. MCT8 staining intensity was similar in IPC expressing low levels of SOX2 or no SOX2 at all (compare **G** and **J**). (**K-N**) Immunostaining of 7-week-old organoids shows enhanced generation of SOX2+/HOPX+ oRGC and a concurrent expansion of the SVZ. Mitotic figures were frequent among the oRGC population in the SVZ (arrows in **N**). (**O-U**) The expanding SVZ was characterized by strong MCT8 expression. Higher magnification views (**R-U**) of the boxed region in **Q** show strong membrane MCT8 staining of HOPX+ oRGC. Dashed lines mark the border between VZ and SVZ in **O** and **P**. Scale bars: 100 µm (**A, B, K, L**), 50 µm (**C, M, O-Q**), 20 µm (**F-J, N, R-U**), 10 µm (**D,E**).

Approximately 1.5 - 2 weeks after initiation of IPC production, RGC in the VZ began to generate HOPX-expressing outer RGC (oRGC), a second type of basal neurogenic progenitors. This *in vitro* timing recapitulates the sequential appearance of IPCs and oRGCs in the human fetal cortex (59). The increased generation of oRGC and their delamination from the VZ resulted in a large expansion of the SVZ between 7 and 8 weeks of culture (**Fig. 4K-M**). Within the SVZ, HOPX+ oRGCs usually assumed more basal positions relative to the TBR2+ IPCs.

Because the generation of an expanded oRGC population is a hallmark distinguishing primate and human cortex development from murine models (44, 60), the observation of a sizeable and highly proliferative oRGC population (see **Fig. 4N**) qualifies our organoids as a viable model to recapitulate human-specific aspects of cortex development. Expression of MCT8 in HOPX+ oRGCs was analyzed on cryosections obtained from 6- to 10-week-old organoids. At all time points analyzed, oRGCs displayed a clear membrane MCT8 staining (**Fig. 4O-U**) and we did not observe appreciable changes in MCT8 staining intensity over time within the oRGC population.

In addition to basal progenitors (IPC and oRGC), the expanding SVZ also inhabits a growing number of newborn neurons generated locally by neurogenic cell divisions of basal progenitors. When addressing TBR1 as one of the first neuronal markers of the excitatory lineage in IF stainings of organoid sections, newborn post-mitotic neurons in the SVZ can be readily identified as TBR1^low^ cells (see **Supplementary Fig. S4**). Since the antibodies available to us did not permit co-staining of MCT8 and TBR1, we assessed MCT8 staining in those SVZ cells expressing none of the NPC markers under the assumption that these cells represent newborn neurons (**Fig. 4)**. Across the depth of the SVZ, we found that newborn neurons displayed a strong MCT8 membrane staining. The MCT8 staining intensity was very similar to that of TBR2+ IPC and we did not recognize subpopulations of SVZ cells characterized by particularly high or low MCT8 staining intensities.

### MCT8 expression in the CP-like zone of cerebral organoids

Neurogenesis of the cerebral cortex proceeds in a sequential inside-out-fashion fashion in that first deep layer neurons are generated while upper layer neurons are produced at later stages (61). This sequence of neuron generation is preserved in cerebral organoids as CTIP2+ deep layer neurons can be observed in the CP-like region as early as 5 weeks of culture whereas the first SATB2+ upper layer neurons do not appear before the seventh week of culture (**Supplementary Fig. S5**). Within ten weeks of organoid culture, the formation of the full scale of neuronal layers is far from complete. The CP-like zone of 10-week-old organoids is characterized by mixed intermingled populations of CTIP2+ and SATB2+ neurons with a variable enrichment of SATB2+ upper layer neurons at the most basal regions (see **Fig. 5A-D**). In 10-week-old organoids, we also noticed that newborn neurons within the SVZ expressed low levels of the deep layer neuron marker CTIP2 but barely express the upper layer neuron marker SATB2 (**Fig. 5G**). Progressively increasing levels of SATB2 staining were detected in deeper regions of the CP (**Fig. 5F**), here often in cells co-expressing CTIP2, whereas strong SATB2 staining was usually restricted to the most basally positioned neurons within the CP-like zone (**Fig. 5E**).

**Figure 5.**
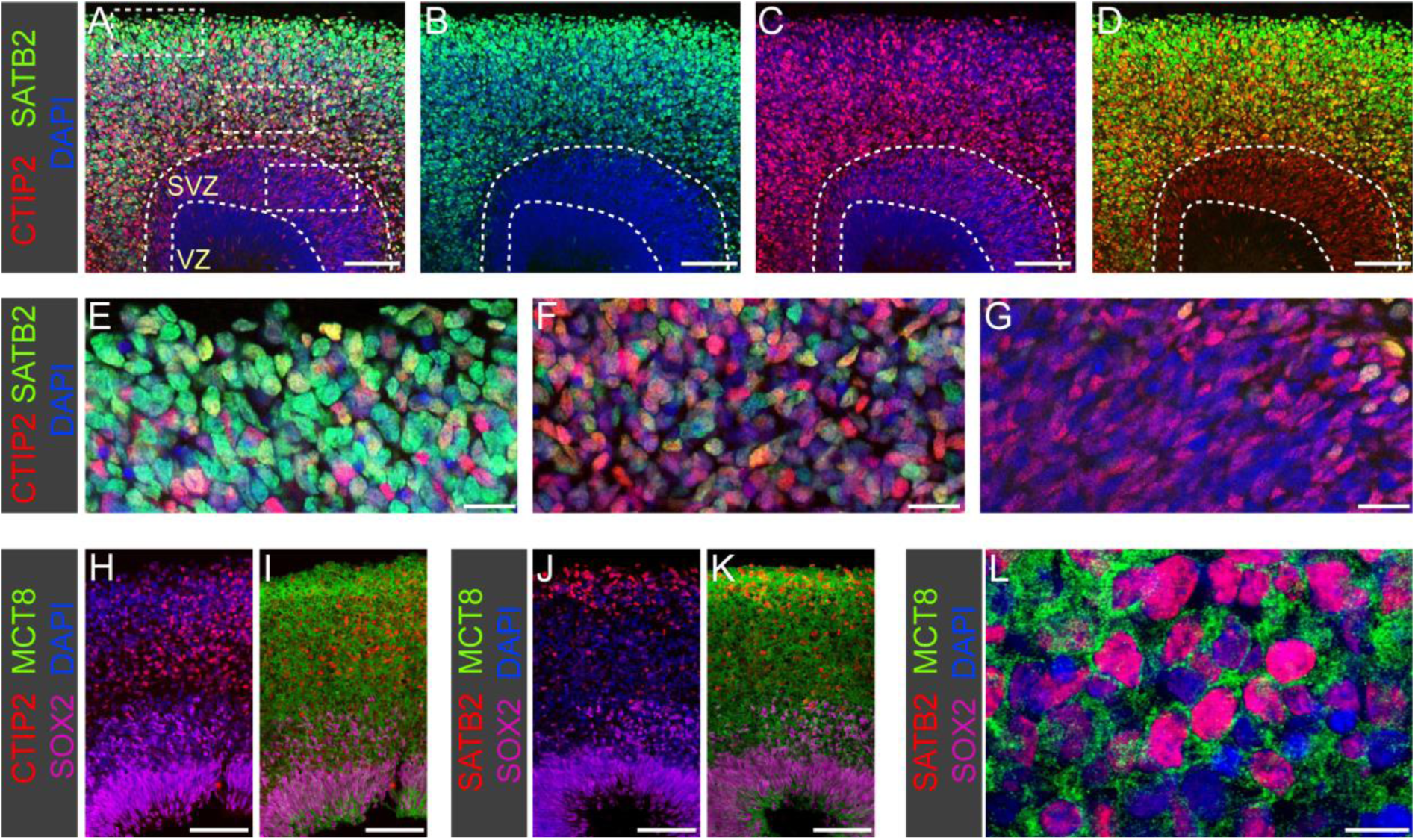
MCT8 expression in neurons. (**A-D**) Immunostaining of 10-week-old organoid for SATB2 (UL neuron marker) and CTIP2 (DL neuron marker) shows that cerebral organoids recapitulate the laminar organization of the CP with early born DL neurons populating inner regions of the CP whereas later born UL neurons are enriched in peripheral regions. Dashed lines mark the border between VZ, SVZ and CP. (**E-G**) High magnification views of the boxed regions highlighted in panel **A**. Note that expression of the UL marker SATB2 is strong in basally positioned neurons (panel **E**), much weaker in deeper CP regions (panel **F**) and almost undetectable in newborn neurons within the SVZ (panel **G**). (**H-K**) In 9-week-old organoids, large field columnar views show abundant MCT8 immunostaining throughout the CP including deep regions populated by DL neurons (**H, I**) and peripheral regions populated by UL neurons (**J, K**). (**L**) Delineation of individual neuronal cell bodies and processes is difficult in the densely populated CP but high magnification views of CP regions confirms MCT8 membrane expression of cortical neurons. Scale bars: 100 µm (**A-D, H-K**), 20 µm (**E-G**), 10 µm (**L**).

A global view of the CP-like zone showed abundant MCT8 staining in this neuronal compartment (**Fig. 5H-K**). In magnified views, membrane MCT8 staining of neuronal cell bodies could be confirmed for both CTIP2+ and SATB2+ neurons (**Fig. 5L**). However, due to the crowding of neuronal cell bodies and the complex and irregular network of axonal and dendritic processes within the CP-like zone, the exact delimitation of cellular boundaries proved difficult in most instances. As a result, it became difficult to conclude on possible differences in the intensity of MCT8 expression between neuronal subpopulations. Collectively, our IF-based analyses of the spatial MCT8 protein expression revealed a predominantly membrane expression of MCT8 in both neuronal progenitors and post-mitotic neurons. Across a culture period from week 2 to week 10 of organoid culture, we observed very limited changes in levels of MCT8 expression.

### Spatial expression of *SLC16A2* and *THRA* mRNA in developing cerebral organoids

We next used the RNAscope technology for fluorescent *in situ* hybridization (FISH) in order to characterize spatiotemporal mRNA expression patterns of *SLC16A2* (encoding MCT8) and *THRA* (encoding TRα) in developing organoids (**Fig. 6**). The *THRA* RNAscope probe mix used in our experiments did not allow to distinguish the expression of the two predominant *THRA* mRNA isoforms, *THRA1* and *THRA2*, expressed in brain tissue (62, 63). All RNAscope staining experiments were combined with IF staining of the NPC marker SOX2 or the neuronal marker TBR1 in order to delineate the border between the VZ, SVZ and CP-like regions. Positive and negative controls were run along each RNAscope experiment.

**Figure 6.**
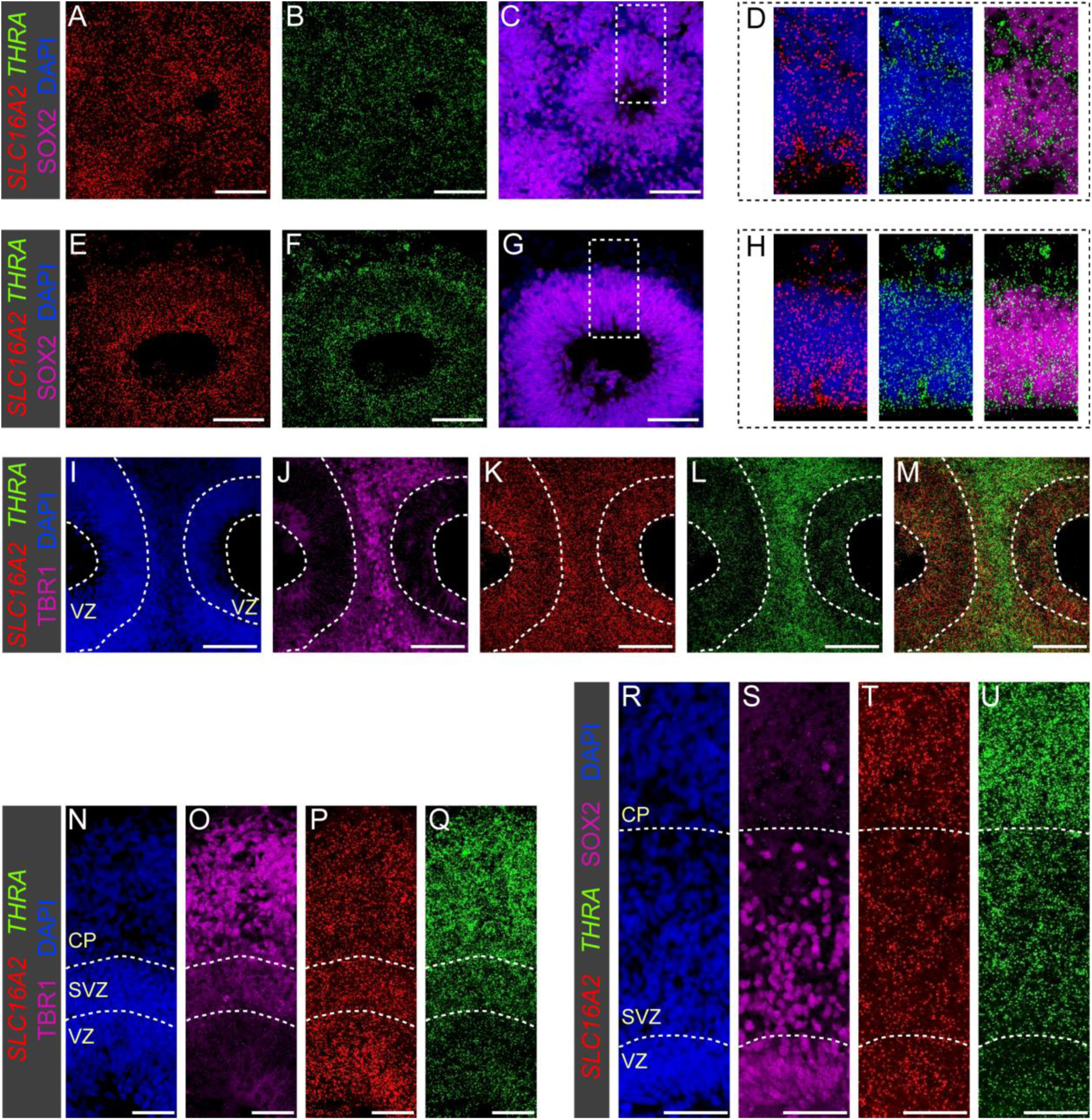
RNAscope analysis of *SLC16A2* and *THRA* mRNA expression in organoids. (**A-D**) Fluorescent *in situ* hybridization of 2.5-week-old organoids shows broad expression of *SLC16A2* and *THRA* mRNA in progenitor cells labeled by SOX2. Panel **D** shows magnified views of the boxed region highlighted in panel **C**. Note that *SLC16A2* and *THRA* expression is also detected in the few SOX2-negative cells (presumably earliest born neurons). (**E-D**) Rosettes of 3.5-week-old organoids showed enriched RNAscope signals for both transcripts in the apical region. *THRA* mRNA also showed enhanced expression in some cells at or near the basal border of the neuroepithelium (see **E** and **H**). (**I-M**) Confocal images of the region between neighboring rosettes in a 5-week-old organoid show similar levels of *SLC16A2* mRNA (**K**) expression in rosettes and adjacent TBR1+ (**J**) neuronal cell layer. RNAscope signals for *THRA* mRNA (**L**) are strongly increased in TBR1+ neurons compared to cell residing in the VZ. Dashed lines mark the apical and basal borders of the VZ. (**N-U**) In organoids cultured for 7 (**N-Q**) and 9 weeks (**R-U**), RNAscope signal intensity for *SLC16A2* mRNA (**P, T**) was comparable across VZ, SVZ and CP regions. In contrast, spatial *THRA* mRNA expression (**Q, U**) was characterized by a gradient from low expression levels in the VZ, intermediate levels in the SVZ and highest expression in neurons of the CP. Dashed lines mark the border between VZ, SVZ and CP in panels **R-U**. Scale bars: 100 µm (**I-M**), 50 µm (**A-C, E-G, N-U**).

In 2.5-week-old organoids that are composed predominantly of SOX2+ NPC, RNAscope signals for *SLC16A2* mRNA were detected with a fairly even distribution across organoid sections (**Fig. 6A**). We also detected RNAscope signals for *THRA* mRNA expression in NPCs of 2.5-week-old organoids confirming an early onset of *THRA* mRNA expression in cerebral organoids. By 3.5 weeks of culture, we noticed spatially enhanced RNAscope signals for *SLC16A2* mRNA near the apical pole of the VZ (**Fig. 6E**). For the sparse population of newborn neurons positioned basal to the VZ in week 3.5 organoids, we observed a positive RNAscope staining but a weak signal intensity (**Fig. 6H**). Similar to *SLC16A2* mRNA, we also noticed an enhanced RNAscope signal intensity for *THRA* mRNA expression near the apical pole of the VZ (**Fig. 6F**). In addition, several cells located immediately basal to the VZ showed a distinct increase in *THRA* mRNA expression (**Fig. 6F**). Based on their specific position adjacent to the VZ, we inferred that these cells with increased *THRA* mRNA expression are most likely early IPC. Enhanced RNAscope signals for *THRA* mRNA were also detected for some of the more basally positioned newborn neurons (**Fig. 6H**).

Along with the generation of a laminar cytoarchitecture by 5 weeks of culture, a strong spatial gradient of *THRA* mRNA expression became apparent in individual cortical units (**Fig. 6I-M**). This spatial *THRA* expression pattern was characterized by low but detectable *THRA* expression levels in SOX2+ NPC within the VZ and high expression levels in neurons of the developing CP-like zone (see **Fig. 6L**). We also noticed that organoids at this stage no longer displayed the enhanced RNAscope signals for *THRA* mRNA near the apical pole of the VZ as seen at earlier stages (see **Fig. 6L, M**). RNAscope analyses of later stage organoids (week 7 and week 9) showed that global *THRA* mRNA expression increased from low levels in the VZ to medium levels in the SVZ and highest levels in the CP layer (**Fig. 6N-U**). In addition, we found that the SVZ displayed the highest heterogeneity of cellular *THRA* mRNA levels (see **Fig. 6U**), presumably due to the mixed cell type composition of this layer, including IPC, oRGC and newborn neurons.

Compared to the marked apico-basal gradient in *THRA* expression, spatial expression of *SLC16A2* mRNA across cell layers of later stage organoids appeared much more uniform. One notable exception was that organoids sampled between 5 and 7 weeks of culture still showed variable levels of *SLC16A2* RNAscope signal enhancement in apical *versus* basal regions of the VZ (see **Fig. 6K, P**). Overall, our RNAscope analyses of *SLC16A2* mRNA expression in later stage organoids (see **Fig. 6N-U**) were consistent with results from MCT8 IF stainings in that both MCT8 protein and *SLC16A2* mRNA were found to be broadly expressed across VZ, SVZ and CP layers.

### T3 treatment of organoids up-regulates expression of known TH-responsive genes

The broad expression of the TH transporter MCT8 and mRNA encoding for TRα implicates a capacity of cerebral organoids to respond to exogenous T3 treatment. To address the responsiveness of *in vitro* grown organoids to an acute T3 treatment, we incubated 44 day-old organoids for 48 h in standard culture media containing a nominal concentrations of 3 nM T3 (basal medium control group) or 50 nM T3 (T3 treatment group) and analyzed bulk mRNA expression of selected genes by RT-qPCR. As shown in **Fig. 7**, a 48 h treatment of organoids with 50 nM T3 induced a strong up-regulation of known T3-responsive genes including *KLF9*, *DBP*, *FLYWCH2* and *CADM2*. In contrast, expression levels of *SLC16A2* and *THRA* mRNA were not affected by this T3 treatment. T3 treatment also did not affect expression levels of cell type markers *SOX2* (RGC), *EOMES* (IPC) and *MAP2* (pan-neuronal).

**Figure 7.**
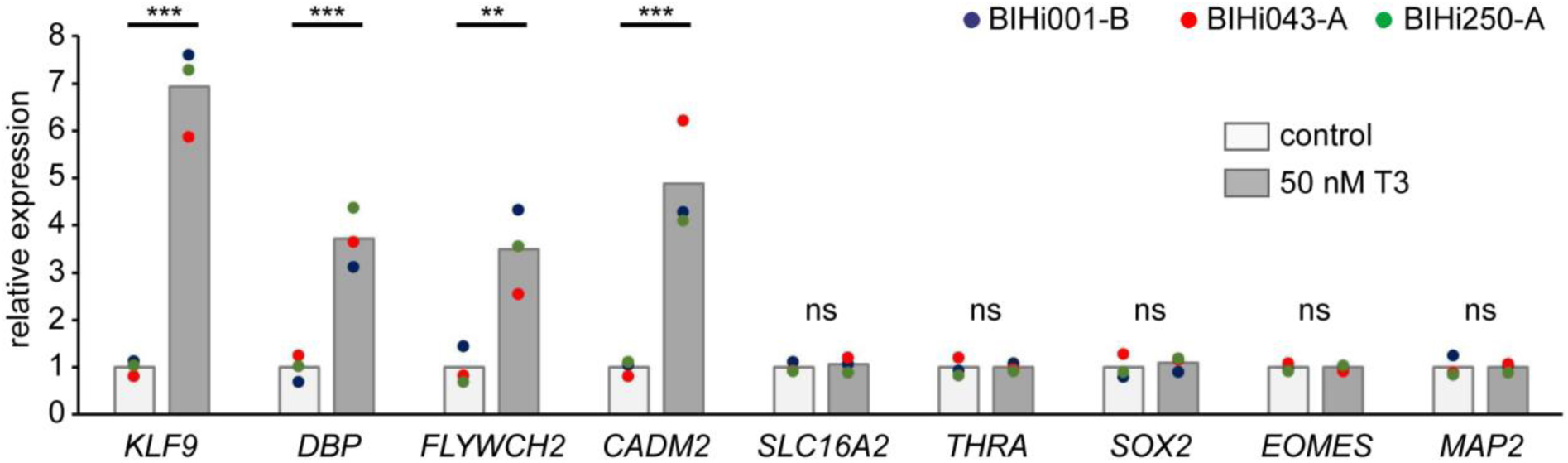
Real-time qPCR quantification of T3 effects on gene expression. 44-day-old organoids derived from three individual hiPSC lines were treated for 48 h with 50 nM T3. Total RNA was isolated from pools of 2-3 organoids and effects of T3 treatment on expression of known T3 response genes (*KLF9, DBP, FLYWCH2, CADM2*), *SLC16A2* and *THRA* as well as cell type markers (*SOX2, EOMES, MAP2*) were determined by RT-qPCR. Colored dots represent mean expression values determined for organoids of individual hiPSC lines. Fold changes relative to the mean of the control group are shown. Unpaired t-test was performed to detect significant differences between treatments using mean values determined for individual hiPSC lines as statistical units (N=3). Asterisks denote significant differences between treatment means. ***, *P*<0.001, **, *P*<0.01. ns, not significant.

## Discussion

Recently, hCOs emerged as powerful three-dimensional *in vitro* models recapitulating key aspects of early human cerebral cortex development. Improved protocols for hCO generation paved the way for new disease modelling approaches and enable multimodal analyses of perturbed neurogenesis resulting from manipulation of specific signalling pathways or gene function (35, 37). These prospects prompted us to explore the value of these model systems for studies on TH function during early human brain development. In this study, we generated hCOs based on a strategy promoting dorsal forebrain tissue generation (46), and verified the successful differentiation of dorsal telencephalic tissue under our culture conditions. In addition, comprehensive immunostaining analyses of organoid tissues collected at various time points across several long-term cultures showed that hCOs indeed displayed hallmarks of human-specific cortex development. Notably, the *in vitro* differentiation from a neuroepithelial stage in 2-week-old hCOs to the appearance of an abundant SATB2+ upper layer neuron population in 10-week-old hCOs took place over a course of 8 weeks. The equivalent developmental processes occur *in vivo* between GW6 and GW14 (49), a similar 8 weeks long time period. Thus, the protracted rate of human cortical neurogenesis is conserved in hiPSC-derived hCOs cultured in our studies.

In addition, hCOs recapitulated the timing of the sequential appearance of specific progenitor populations seen in human fetal cortex and in particular the generation of abundant HOPX+ oRGC. The latter finding reflects an important human-specific feature of hCOs since oRGC are rare in the developing rodent cortex (60, 64). oRGC are equipped with self-renewal capacity and a clonal output that is 10-times higher compared with mouse RGC (60). The abundant oRGC population in the human SVZ therefore creates a second major germinal region contributing to the size expansion of the human gyrencephalic cortex (31). Similar to the different progenitor cell populations, we also verified that hCOs recapitulate the sequential generation of post-mitotic neurons seen *in vivo* including the generation of early TBR1+ neurons followed by generation of TBR1+/CTIP2+ deep layer neurons and finally late-appearing SATB2+ upper layer neurons (61). Moreover, 10-week-old organoids also displayed an inside-out pattern of neurogenesis in that later generated SATB2+ neurons migrated progressively beyond early born deep layer neurons to populate more superficial positions (65).

hCOs cultured according to our protocol for up to 10 weeks did not contain astrocytes, oligodendrocytes or interneurons (data not shown). While astrocytes and oligodendrocytes are generated from the same initial neuronal stem cell pool that drives cortical neurogenesis, the absence of astrocytes and oligodendrocytes in our hCOs can be explained by the delayed onset of glia generation occurring in the second trimester of human pregnancy (66, 67). The absence of cortical interneurons in our hCOs is related to their different place of origin, the ganglionic eminences of the ventral forebrain from where interneurons migrate dorsally into the developing neocortex (68). Collectively, our analysis of neurodevelopmental markers suggest that the molecular neuroanatomy of hCOs generated by our protocol could provide a unique opportunity to address regulatory functions of TH in a complex 3D models recapitulating cortical wall morphogenesis happening *in vivo*.

To date, information about developmental expression profiles of critical regulators of TH action is still very sparse with respect to early human cortex development (10, 57). To fill this gap, we here analyzed expression of a critical TH transporter, MCT8, on protein and mRNA levels as well as expression of *THRA* mRNA in hCOs. MCT8 immunostainings showed membrane staining of all major cell types including neuronal progenitors (RGC, IPC and oRGC) as well as post-mitotic neurons, and high-resolution confocal imaging showed if any only minor differences in staining intensities. A similar broad distribution of MCT8 protein has previously been reported in cortex tissue sampled from fetuses at GW14 and later stages (10, 57). Our orthogonal RNAscope analyses of hCO tissue corroborated a broad expression of *SLC16A2* mRNA across all cell layers.

One notable exception, however, was an enhanced MCT8 immunostaining of the most apical portion of the VZ lining the lumen. A recent study similarly observed a more prominent immunostaining intensity at the apical neuroepithelial surface of a GW20 fetal cortex sample (10). The recurrent observation of enhanced MCT8 protein expression at the apical surface lining the luminal space both *in vivo* and *in vitro* suggest two things. First, MCT8 might play an important role for TH uptake from the cerebrospinal fluid into RGC highlighting the relevance of this border for TH supply of the developing cortex. Second, conservation of this local accumulation of MCT8 protein in hCOs indicates that MCT8 distribution in RGC is under intrinsic cellular control likely related to the overall apico-basal polarity that RGCs display both *in vivo* and *in vitro*. When combining results from the current hCO study and several previous analyses of fetal cortex tissues (10, 11, 57), one can conclude that cortex development is characterized by broad MCT8 expression from early neuroepithelial stages throughout mid-gestation neurogenesis.

In contrast to the limited changes in spatiotemporal MCT8 expression, our RNAscope analyses revealed a much more dynamic developmental profile for *THRA* mRNA expression during hCO differentiation. The demonstration of *THRA* mRNA expression in the neuroepithelium of early-stage hCOs is of special interest as it implicates a competence of early neuronal progenitor cell populations to respond to TH, particular given the demonstrated concurrent expression of MCT8 protein. The developmental stage of the *in vitro* neuroepithelium of 2.5-week-old hCOs corresponds to human fetal cortical tissue at GW6 to GW7. Our observations at the mRNA level are in line with previous PCR-based studies reporting *THRA* mRNA expression in human fetal cortex tissue at GW7 to GW8 (69, 70).

Unavailability of specific TRα antibodies limited our analyses so far to the determination of mRNA expression and it remains unknown if relevant quantities of TRα protein are synthesized during these early developmental stages. This early neuroepithelium is the precursor structure of the cortical VZ containing RGC as the major neuronal stem cell population. The observation that, at least in rodent experimental studies, maternal hypothyroidism has been linked to perturbed RGC function (71–73) warrants further studies to verify the actual onset of functional TRα protein expression in the VZ.

Another key finding of our study was the marked up-regulation of *THRA* mRNA expression along the neuronal differentiation trajectory. This was particularly evident when mapping RNAscope signals for *THRA* mRNA on the layered cytoarchitecture of hCOs at later stages. While low levels of *THRA* mRNA were maintained in the VZ of older organoids, *THRA* mRNA expression was gradually increased in the SVZ and highest expression levels were detectable in neurons populating the CP. Rodent data on the spatial *THRA* expression in developing cortex similarly reported highest expression levels in the CP (74–76). However, to our knowledge there are no studies describing spatial *THRA* expression profiles in human fetal cortex at high resolution.

When integrating our observations from different time points, a model emerged that there are only limited temporal changes of *THRA* mRNA expression within a given cell population but that *THRA* mRNA expression increases greatly along the differentiation trajectory from RGC to excitatory neurons. A quantitative confirmation of such a model is difficult based on spatial expression data alone but an integrated analysis of published single cell transcriptome of human fetal cortex tissues (44, 49, 77, 78) holds promise to address this topic with cell type resolution.

In this study, we also performed a first proof-of-principle experiment to document that these millimetre-sized organoids respond sensitively to a short-term exposure to elevated T3 media concentrations, as evident from the robust up-regulation of known T3 response genes (3, 4). The complex tissue composition of hCOs generated by our protocol certainly warrants the use of single cell RNA sequencing methods to capture eventually TH action in a more comprehensive and cell type-specific manner. An important outcome of this still limited gene expression analysis was the observation of very homogenous responses of hCOs derived from different hiPSC lines. This finding supports our view that implementation of quality control measure promotes the reproducibility of organoid cultures.

Although hiPSC became a key tool for modelling human neuronal development and disorders, differences in the propensity of individual hiPSC lines to differentiate into the desired cell lineages remains a major source of variability (79). While most of the data of this study have been generated for hCOs derived from the male BIHi001-B line, additional experiments with two other hiPSC lines (BIHi043-A and BIHi250A) confirmed the robustness of our protocol. Efficiency of dorsal telencephalic patterning, propensity to form of multi-layered structures and the timing of key developmental processes related to cerebral cortex development were very similar across organoid cultures using these three hiPSC lines. However, similar to previous reports (43, 44), we observed examples of inter-experimental differences in the efficiency of cortical specification (e.g., when using line BIHi005-A), sometimes even when using different batches of the same hiPSC line. To permit the identification of organoid batches that failed cortical lineage specification already at early stages of the culture protocol, we devised batch quality control measures that include monitoring of gross morphological characteristics and most importantly, a rigorous verification of dorsal forebrain specification during the fourth week of culture. Early identification and termination of organoid batches that are non-compliant with quality criteria helps greatly to reduce time and costs given the long duration of organoid culture protocols of up to 10 weeks. In addition, we argue that implementation of rigorous quality control assessments is critically required to ensure a “normotypic” timeline of key developmental processes during independent organoid cultures. This information will be of great importance to efficiently guide the timing of experimental interventions and sampling regimes in forthcoming studies exploring the consequences of altered TH action on cerebral development.

In conclusion, this study showed that hiPSC-derived human cerebral organoids recapitulate key aspects of human first trimester cortical development and provide a unique platform to examine the critical role of TH action for early human brain development. Additionally, we highlighted the importance of the quality control measures to ensure robust and reproducible endpoint measurements related to TH signalling. The information that we obtained about temporal progression of neurogenesis in organoids and the concurrent expression of MCT8 and THRA provides critical guidance for hCO culture experiments aiming at a comprehensive interrogation of TH-responsive gene expression profiles using high resolution techniques such as single cell RNA sequencing.

## Author Contribution

A.G. processed all organoid samples, performed sectioning, established all immunofluorescence assays, performed immunofluorescence staining and RNAscope assays, prepared figures, and wrote the article together with R.O. A.A.J.B. performed sectioning, immunofluorescence staining and RNAscope assays. V.F.V. established the organoid culture protocol and performed organoid cultures and qPCR assays. M.M. performed organoid cultures. H.S. provided all hiPSC lines and equipment for hiPSC maintenance and organoid cultures. R.O. performed confocal microscopy. P.K. and R.O. conceived and supervised the study. All authors reviewed and edited the manuscript.

## Acknowledgments

We thank J. Küchler for excellent technical support in organoid culture experiments.

## Additional Information

Supplementary information accompanies this paper.

