## Supplemental Figures 1-5 for "Spatiotemporal expression of thyroid hormone transporter MCT8 and *THRA* mRNA in human cerebral organoids recapitulating first trimester cortex development"

**Supplementary Figures S1 – S5**

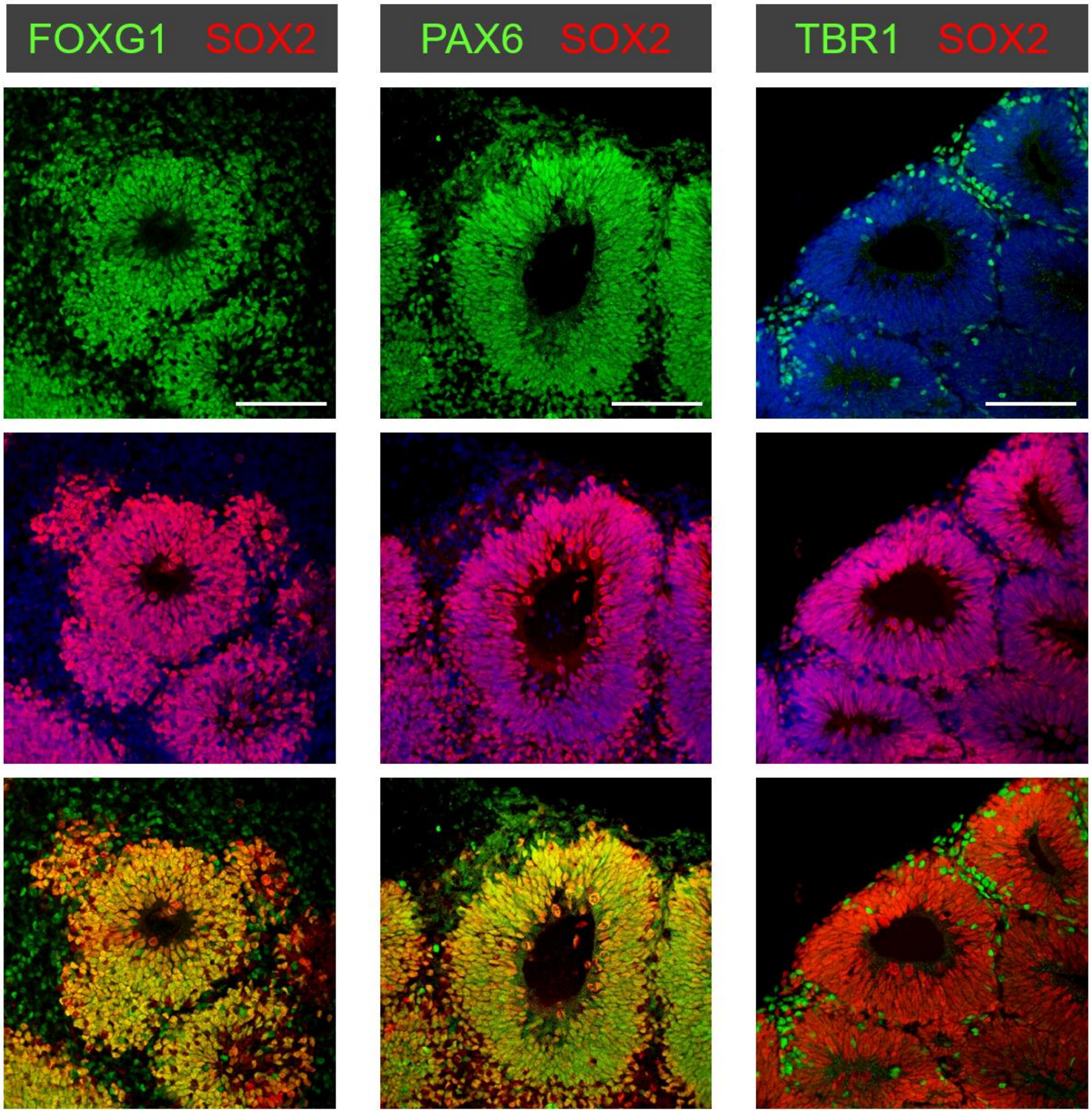


**Figure S1**. Quality control assessment of dorsal telencephalic patterning during the fourth week of organoid culture (quality checkpoint #3). Correct patterning of early stage organoids was verified by expression of FOXG1 and PAX6 in SOX2+ neuronal progenitors and expression of TBR1 in newborn neurons. For this critical quality checkpoint #3 of our culture protocol, a subset of organoids was analyzed between culture day 24 and day 28 of any given culture experiment. Images show results of immunostaining of 24-day-old organoids derived from the BIHi001-B line. Scale bars: 100 µm.

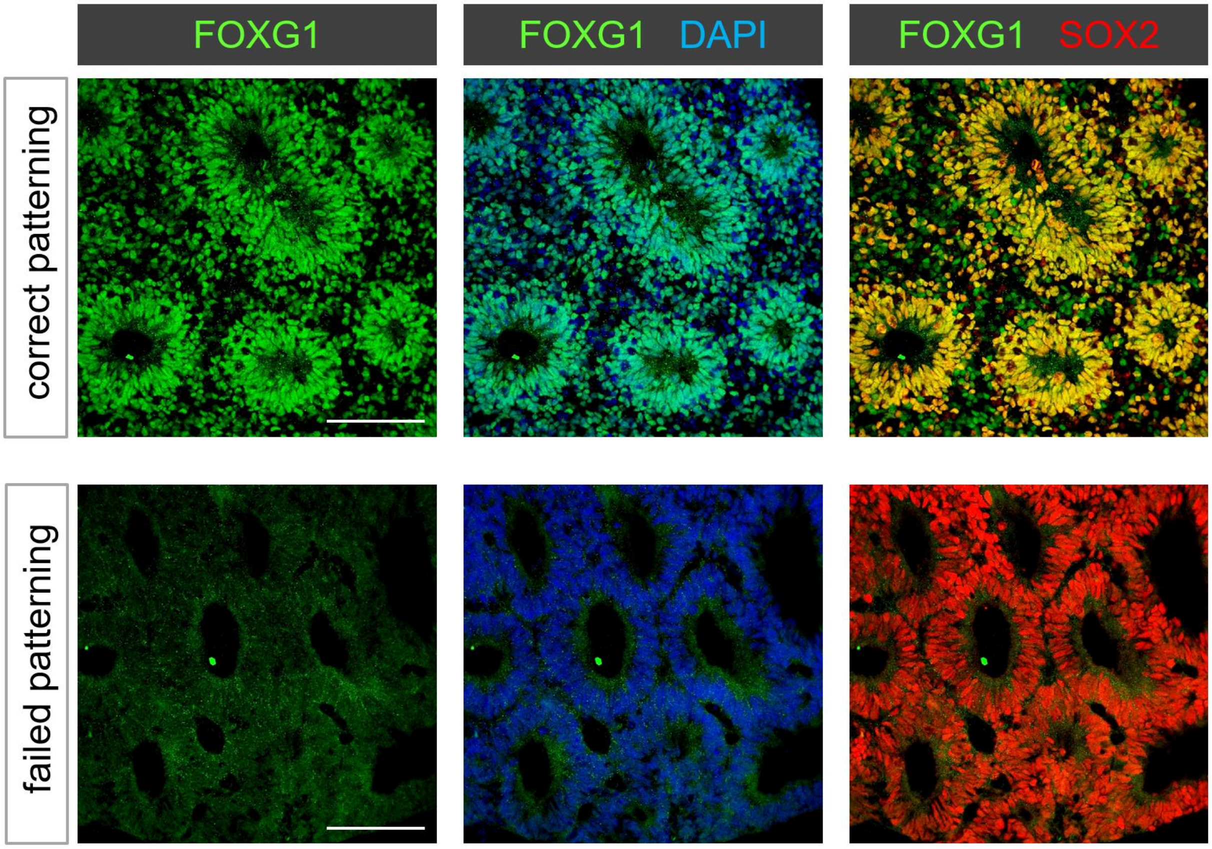


**Figure S2**. Examples of organoid batches that passed (upper panel) or failed to pass (lower panel) criteria of quality checkpoint #3 (assessment of dorsal telencephalic patterning). Upper panel shows positive FOXG1 immunostaining of 27-day-old organoid derived from BIHi250-A line. Lower panel shows lack of FOXG1 expression in 27-day-old organoid derived from BIHi005-A line. Note that organization of SOX2+ cells into rosette-like structures is not a sufficient criterion to verify successful generation of cerebral cortical tissue. Organoids from the non-compliant BIHi005-A batch were initially compliant with gross morphological criteria described for quality checkpoints #1 (EB formation) and #2 (neuroepithelium). Failed cortical lineage specification was only revealed by analysis of marker expression in early-stage organoids. Scale bars: 100 µm.

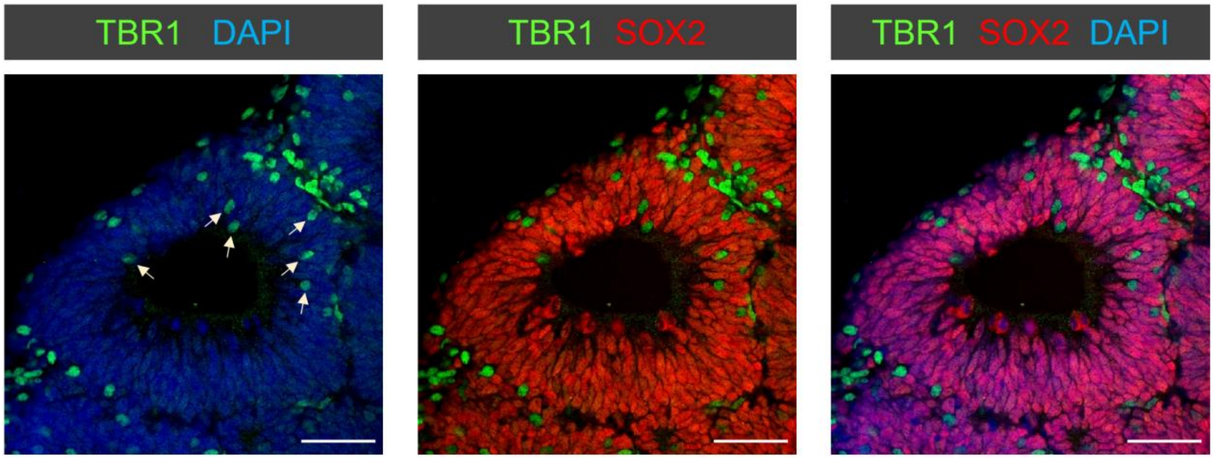


**Figure S3**. Direct neurogenesis in early-stage organoids. Distribution of TBR1+ neurons (arrows) in ventricular zone of 3.5-week-old organoids. The distribution profile of TBR1+ cells indicates that RGC divisions in the VZ produce daughter cells that directly differentiate into neurons bypassing the generation of basal neurogenic progenitors such as IPC and oRGC. When judged from prevalence of TBR1+ cells within the VZ, direct neurogenesis was prominent in organoids during culture weeks 3 and 4 and declined at later stages. Scale bars: 50 µm.

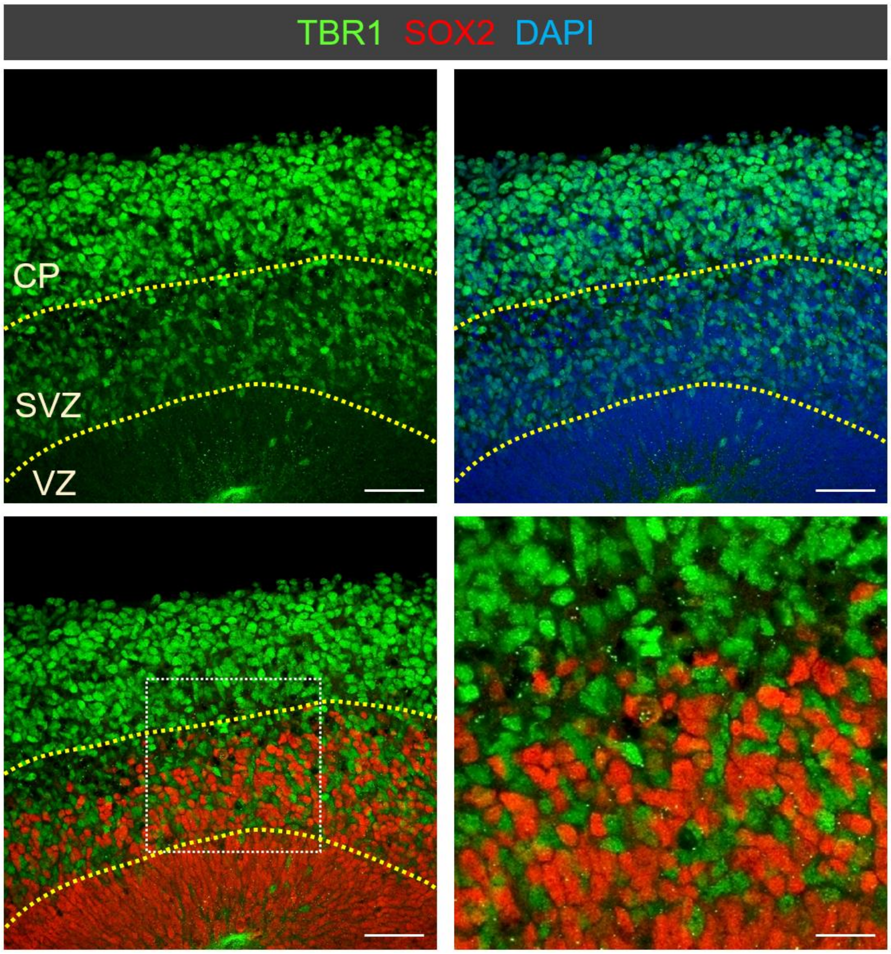


**Figure S4**. Newborn TBR1+ neurons in the SVZ of 7-week-old organoids are characterized by a lower immunostaining signal intensity compared to more mature TBR1+ neurons that migrated out to the CP. Lower right image shows a magnified view of the boxed region marked in the lower left image. Dashed lines mark the border between VZ, SVZ and CP. Scale bars: 50 µm (20 µm in lower right magnified view image).

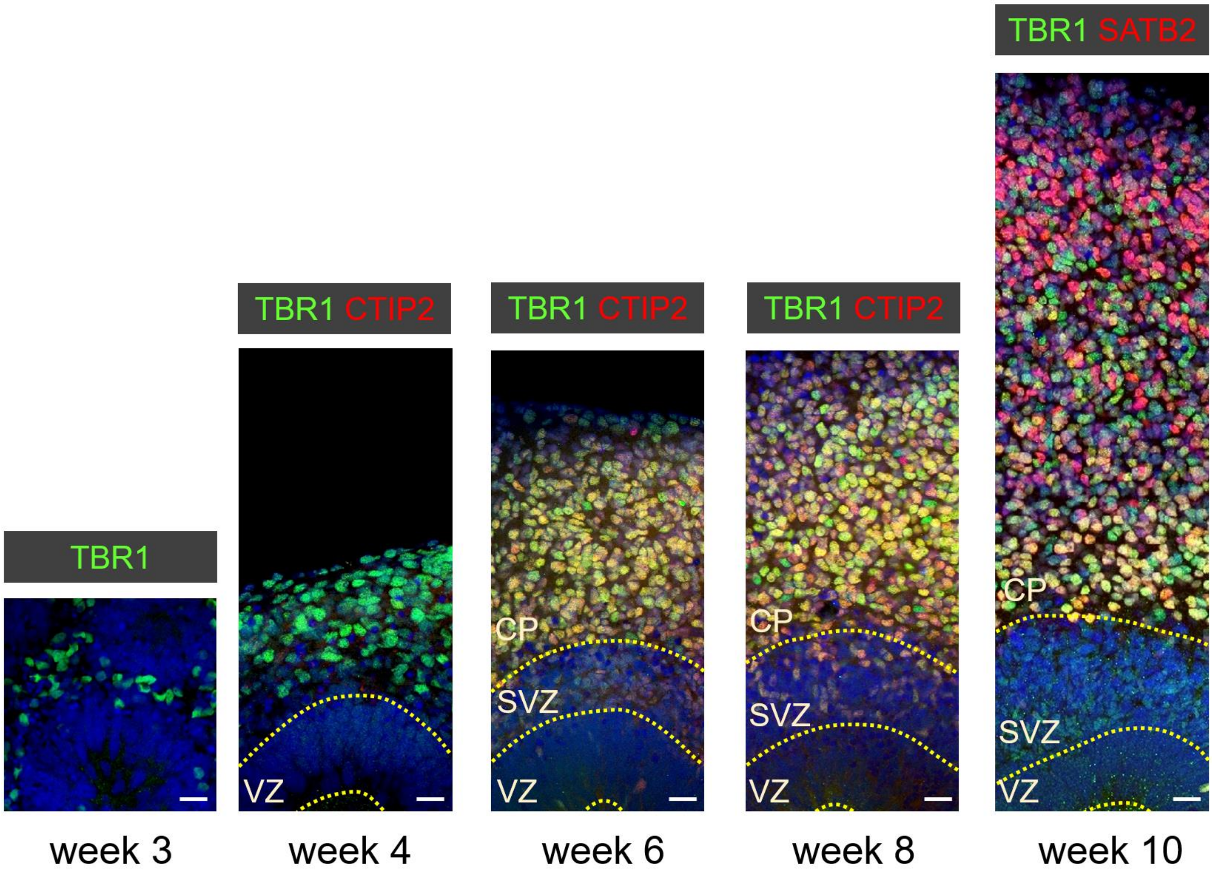


**Figure S5**. Sequential generation of a diverse repertoire of cortical neurons during a 10-week organoid culture recapitulates the inside-out pattern of neuronal layering. A first wave of neurogenesis between culture weeks 2 and 4 produces mainly TBR1+ neurons that do not yet express deep layer neuron markers such as CTIP2. A next wave of neurogenesis (4 to 8 weeks of culture) produces neuronal subtypes that will eventually populate the deepest layers of the cortical wall. These deep layer neurons express high levels of CTIP2. As corticogenesis proceeds further, newly generated neurons migrate radially throughout the deep neuronal layer to occupy successively the upper layers of the cortical plate. Accordingly, first expression of the upper layer neuron marker SATB2 becomes detectable during the 8th week of culture and a prominent SATB2+ neuronal layer is visible in the upper region of the cortical plate in 10-week-old organoids. Dashed lines mark the border between the ventricular zone (VZ), subventricular zone (SVZ) and cortical plate (CP). Scale bars: 20 µm.
